# The PhytoClust Tool for Metabolic Gene Clusters Discovery in Plant Genomes

**DOI:** 10.1101/079343

**Authors:** Nadine Töpfer, Lisa-Maria Fuchs, Asaph Aharoni

**Affiliations:** Department of Plant Science and the Environment, Weizmann Institute of Science, Rehovot, Israel; Hochschule für Wissenschaft und Technik, Berlin, Germany

## Abstract

The existence of Metabolic Gene Clusters (MGCs) in plant genomes has recently raised increased interest. Thus far, MGCs were commonly identified for pathways of specialized metabolism, mostly those associated with terpene type products. For efficient identification of novel MGCs computational approaches are essential. Here we present PhytoClust; a tool for the detection of candidate MGCs in plant genomes. The algorithm employs a collection of enzyme families related to plant specialized metabolism, translated into hidden Markov models, to mine given genome sequences for physically co-localized metabolic enzymes. Our tool accurately identifies previously characterized plant MBCs. An exhaustive search of 31 plant genomes detected 1232 and 5531 putative gene cluster types and candidates, respectively. Clustering analysis of putative MGCs types by species reflected plant taxonomy. Furthermore, enrichment analysis revealed taxa- and species-specific enrichment of certain enzyme families in MGCs. When operating through our web-interface, PhytoClust users can mine a genome either based on a list of known cluster types or by defining new cluster rules. Moreover, for selected plant species, the output can be complemented by co-expression analysis. Altogether, we envisage PhytoClust to enhance novel MGCs discovery which will in turn impact the exploration of plant metabolism.

## Introduction

Operon-like features known as Metabolic Gene Clusters (MGCs) are commonly found in fungal genomes. They represent genes of a particular metabolic pathway that are physically linked in the genome. It was long assumed that MGCs occur as an exception in plants (1). However, this assumption has been challenged in recent years; more than 20 MGCs have been experimentally characterized in a variety of species, the majority associated with plant specialized metabolism (2–6). Plant MGCs span genomes of both mono- and dicotyledonous species mediating the biosynthesis of end-products associated with different chemical classes including benzoxazinoids, cyanogenic glycosides, terpenoids, and alkaloids (3). Significantly, the range of reported plant MGCs includes biosynthetic reactions for the synthesis of pharmaceutically and agronomically important chemicals, *e.g.,* the anti-tumor alkaloid noscapine in poppy, the anti-nutritional steroidal alkaloids in potato or bitter cucurbitacin triterpenoids in cucumber (7–9).

A common descriptor of plant MGCs is the adjacent localization of at least three, occasionally two (10), non-homologous biosynthetic genes that encode enzymes involved in the synthesis of specialized metabolites (4). It appears that plant MGCs evolved independently from prokaryotic operons. Typically, plant MGCs are composed of one gene encoding a so-called “signature enzyme”, *i.e.* an enzyme which catalyzes the first committed step of the biosynthetic pathway and synthesizes the scaffold of the following specialized metabolites. The remaining genes encode subsequent “tailoring enzymes” which modify the scaffold to form the desired end-product (4). Moreover, as signature genes share homology with genes of plant primary metabolism it is widely assumed that plant MGCs were formed by the recruitment of additional tailoring enzymes through gene duplication and neo-functionalization.

The finding of MGCs in plant genomes accelerates pathway discovery and the capacity to metabolically engineer desired specialized metabolites (11, 12). At the same time, it also increases the demand for plant genome sequencing and genome-mining. Consequently, several *in silico* approaches were undertaken to systematically screen plant genomes for putative MGCs. In one of the earliest attempts the authors presented co-expression analysis of neighboring genes in Arabidopsis across 1469 experimental conditions (13). Based on their analysis 100 putative clusters were identified of which 34 were significantly co-expressed, containing 3 to 22 genes and 27 duplicated gene pairs.

As part of a more extensive analysis that investigated diversification of metabolism in 16 plant species, Chae *et al.* (2014) examined the clustering of metabolic genes across primary and specialized metabolism. MGCs were detected based on the following three criteria: (i) all genes in a cluster must be associated with a four-part EC number, (ii) more than one distinct EC number must be represented in the cluster, and (iii) all genes in a cluster must be contiguously located on the same chromosome. The authors found approximately one-third of the metabolic genes in Arabidopsis, soybean, and sorghum, and one-fifth of the genes in rice matching these criteria. In the case of Arabidopsis, the authors tested whether clustered genes exhibit significant co-expression patterns. The results indicated that putative MGCs associated with specialized metabolism are more likely to be co-expressed than their putative counterparts from primary metabolism. However, using the same data set, Omranian *et al.* (2015) demonstrated that conclusions drawn from these results should be handled with care as patterns of co-expression between specialized and non-specialized metabolic pathways may differ depending on the considered correlation.

A different study focused merely on the large class of terpenoid specialized metabolites by examining the distribution of pairs of terpene synthases (TS) and cytochrome p450s (CYPs) across 17 sequenced plant genomes (15). TS and CYPs have been found in several reported MGCs as the product of TS activity is typically further decorated by CYPs enzymes. It appeared that physically co-localized pairs of TS/CYPs were found much more frequently than by chance. By further investigating the predominance of certain TS/CYPs pairs across species the authors uncovered different mechanisms of pathway assembly in eudicots and monocots.

AntiSmash, the primary tool for online MGCs search was initially developed for the detection of clusters in fungi and bacteria (16–18). The tool employs a hidden Markov model (HMM) search algorithm to detect genes that are specific for certain types of known clusters *i.e.* the tool is confined to already characterized gene cluster types.

Despite the insights generated by these studies, a generalized framework for identifying plant MGCs is required. Such an approach should be applicable to a large set of genomes as well as be comprehensive and comprise searching capabilities beyond the known MGCs types. Ideally, it should combine sequence-based information with genomic outputs such as gene co-expression. Following the needs described above we developed and applied PhytoClust, an *in silico* MGC prediction tool. PhytoClust allows the search for known plant MGC types as well as mining for novel types of clusters (*i.e.* in terms of enzyme class composition). Co-expression analysis of genes located in candidate clusters is available for selected plant species. We anticipate that PhytoClust will enhance the characterization of novel MGCs in a wide range of plant species as new genome assemblies become available.

## RESULTS

### The PhytoClust tool to detect metabolic gene clusters in plants

We developed PhytoClust, a software that uses a collection of enzyme families of plant specialized metabolism (Table 1) translated into hidden Markov models (HMMs) and mines a given genome sequence for co-localized metabolic enzymes of these families (“marker enzymes”) (Supplementary Information 1). Using the core implementation of AntiSmash (18) the search query parameters are defined by “cluster rules” which include (i) the names of enzyme families of interest, (ii) the span of chromosomal region in which genes are clustered (“cluster range”) and (iii) the flanking region to be searched for additional marker enzymes (Figure 1). PhytoClust enables the search for currently known gene cluster types as well as setting individual criteria with a combination of up to four enzyme families. It also allows searching for tandem repeat gene clusters. Once the search query has been processed the results can be examined in a web-browser or downloaded for further offline analysis. Additionally, for a small collection of well annotated genomes and available transcriptome data sets we developed a co-expression module that provides co-expression analysis for genes located in candidate clusters based on a user-defined co-expression threshold and visualizes the results as heat-maps. PhytoClust is available via a web-server at www.PhytoClust.ac.il.

**Table 1:**
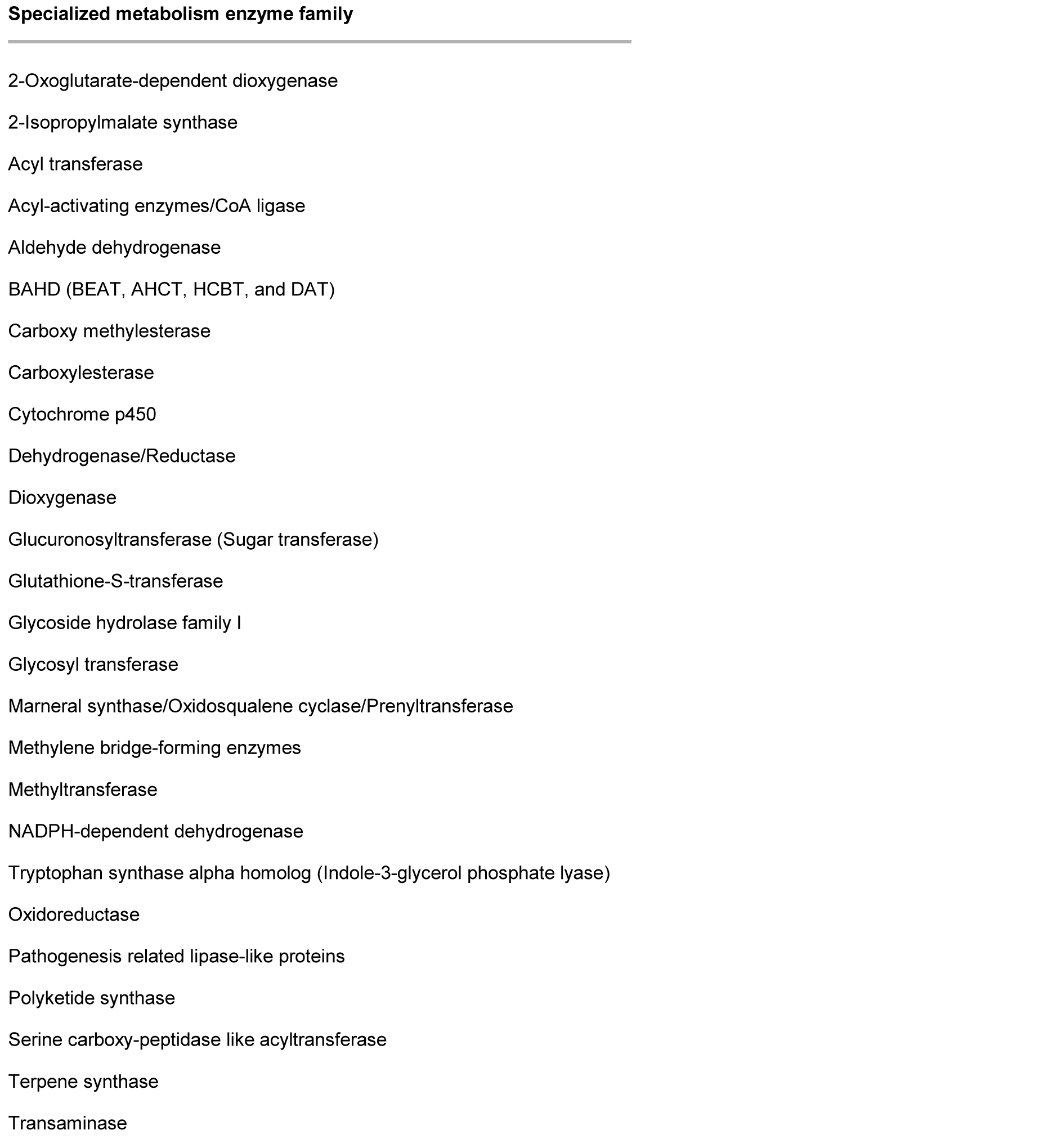
List of enzyme families associated with specialized metabolism. Literature screening and manual curation to avoid redundancy resulted in a collection of enzyme families associated with plant specialized metabolism; see (4, 31–33) and references within.

**Figure 1:**
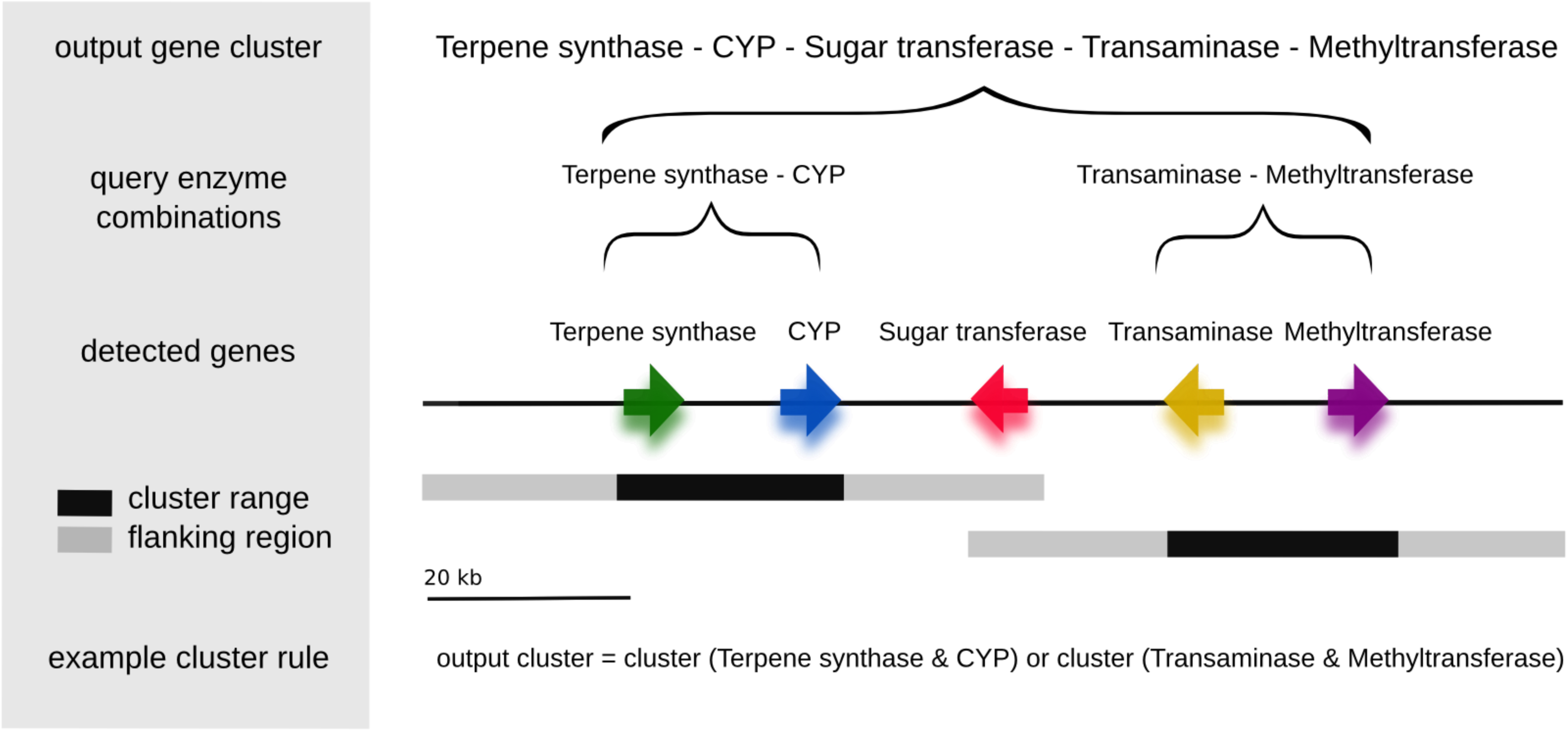
Schematic representation of the cluster detection algorithm. PhytoClust searches a given plant genome for genes representing enzyme families associated with specialized metabolism (“marker enzymes”) based on a hidden Markov model search algorithm. The search query is formalized by “cluster rules” that contain the query enzyme families and parameters for the cluster range (sequence range in which the marker enzymes are to be detected) and the flanking region (proximity of the cluster). A hit is recorded if genes encoding the query enzymes are detected within the cluster range. Additional marker enzymes from the marker enzyme library which are detected within the cluster range or the flanking region of the cluster will be automatically added to the results. Analogously, neighboring gene clusters will be merged if they are within reach of each other’s flanking region.

### PhytoClust accurately predicts known plant metabolic gene clusters

To demonstrate the quality of PhytoClust we first attempted to detect established plant MGCs located in genomes having high-quality assemblies. These included the Arabidopsis marneral (19) and thalianol clusters (2), *Solanum lycopersicum* (tomato) terpene cluster (20), *Solanum tuberosum* (potato) and *Solanum lycopersicum* steroidal glycoalkaloids cluster (8), *Oryza sativa* (rice) momilactone (21) and phytocassane cluster (22), and the *Zea mays* (maize) DIMBOA cluster (23–26). Based on the structure of these MGCs we defined cluster rules for the “known-gene-cluster-search-module” (Supplementary Information 2). Significantly, we accurately find the above mentioned MGCs among few candidate MGCs that fit the search criteria (between one to five MGC candidates, depending on the cluster type and species). The search criteria and results are summarized in Figure 2.

**Figure 2:**
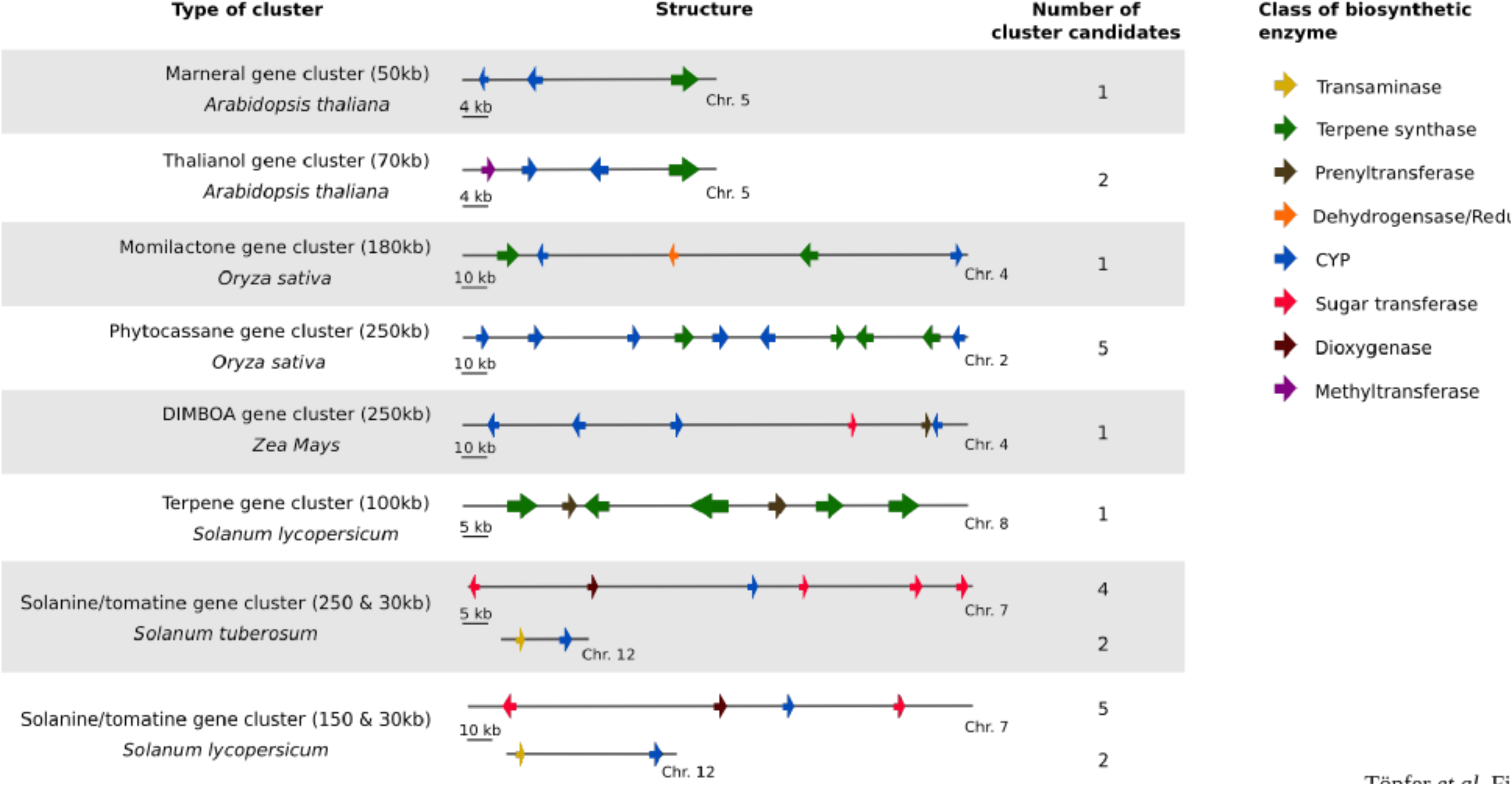
Graphical representation of known MGCs investigated in this study. ”Type of cluster” shows the cluster name, species, and the proximity in which the respective enzymes are located. “Structure” gives a graphical representation of the cluster’s physical map (Approximation). “Number of detected cluster” lists the total number of hits obtained when running PhytoClusts’ “known-gene-cluster-search” including the characterized gene cluster. All eight gene clusters in species with high-quality genome assemblies were detected with high accuracy. Image adapted from (4).

### Benchmarking PhytoClust with existing tools for MGCs identification

To benchmark PhytoClust against existing MGCs prediction tools we compared our results to AntiSmash (version 3.0.5); to our knowledge, the only currently available tool containing information regarding plant specific MGCs. As plant genomes are very large and due to restrictions in upload file size of the AntiSmash server, we were not able to upload the whole plant genome assemblies, but had to limit our test cases to confined sequences within the plant genomes containing the actual MGCs shown in Figure 2. AntiSmash identified the marneral gene cluster in Arabidopsis (detected as Terpene cluster), the momilactone and phytocassane gene cluster in rice (detected as Terpene - Momilactone and biosynthetic gene cluster and Terpene - Phytocassane /Oryzalides biosynthetic gene cluster, respectively), the terpene gene cluster in tomato (detected as Terpene gene cluster) and the chromosome 12-located part of the steroidal glycoalkaloid gene cluster in tomato (detected as a Terpene gene cluster). MGCs not detected by AntiSmash included the thalianol gene cluster in Arabidopsis, the chromosome 7-located part of the glycoalkaloid gene cluster in tomato and potato, the chromosome-12-located part of the steroidal glycoalkaloid gene cluster in potato, and the DIMBOA gene cluster in maize. All of the mentioned clusters were accurately detected by PhytoClust, as outlined in the previous section. The results of the comparison are summarized in Table 2. As our search tool is based on the core-implementation of AntiSmash, we conclude that the observed differences are mainly due to the different search criteria implemented in the cluster rules in PhytoClust (see also Materials and Methods).

**Table 2:** Comparison between PhytoClust and AntiSmash (Version 3.0.5). As a test case, the eight characterized gene clusters in Arabidopsis, rice, tomato, potato, and maize were searched. Note, that Solanine 1, Tomatine 1 refer to the split cluster part located on chromosome (Chr.) 7 and Solanine/Tomatine 2 refer to the split cluster part located on 12. The column “PhytoClust” shows the results obtained using the “known-gene-cluster-search-modus”. Column “AntiSmash” shows the results obtained when applying the AntiSmash pipeline to the same sequence within the respective plant genome.

| Cluster |  | PhytoClust | AntiSmash |
| --- | --- | --- | --- |
| ***Arabidopsis thaliana*** |  |  |  |
| Chr. 5<br>Position: 17003646-17119359 | Marneral | Arabidopsis Marneral | Terpene |
| Position Chr. 5<br>Position: 19414827-19481538 | Thalianol | Arabidopsis Thalianol – Arabidopsis Marneral | / |
| ***Oryza Sativa*** |  |  |  |
| Chr. 4<br>Position: 5298173-5506541 | Momilactone | Oryza S Phytocassane – Oryza S Momilactone | Terpene - Momilactone biosynthetic gene cluster |
| Chr. 2<br>Position: 21607356-21934209 | Phytocassane | Oryza S Phytocassane | Terpene - Phytocassane /Oryzalides biosynthetic gene cluster |
| ***Solanum lycopersicum*** |  |  |  |
| Chr. 8<br>Position: 545618-624754 | Terpene | Solanum L Terpene | Terpene |
| Chr. 7<br>Position: 54353341-54742851 | Tomatine 1 | Solanum L Tomatine 1 |  |
| Chr. 12<br>Position: 788715-1062587 | Solanine/Tomatine 2 | Solanum L Tomatine 1 - Solanum T/L Solanine/Tomatine 2 | Terpene |
| ***Solanum tuberosum*** |  |  |  |
| Chr. 7<br>Position: 41670011-42005447 | Solanine 1 | Solanum T Solanine 1 | / |
| Chr. 12<br>Position: 5803461-5978242 | Solanine/Tomatine 2 | Solanum T solanine 1-Solanum T/L Solanine/Tomatine 2 |  |
| ***Zea mays*** |  |  |  |
| Chr. 4<br>Position: 3001405-3322257 | DIMBOA | Zea M DIMBOA | / |

### PhytoClust identifies putative novel MGCs in plant and algal genomes

To assess the relationship between MGCs and plant taxonomy we applied PhytoClust to a selection of 31 high-quality genome sequences from the green lineage (Figure 3), including three algae species, a moss and a lycophyte, 12 monocotyledonous and 14 dicotyledonous species.

**Figure 3:**
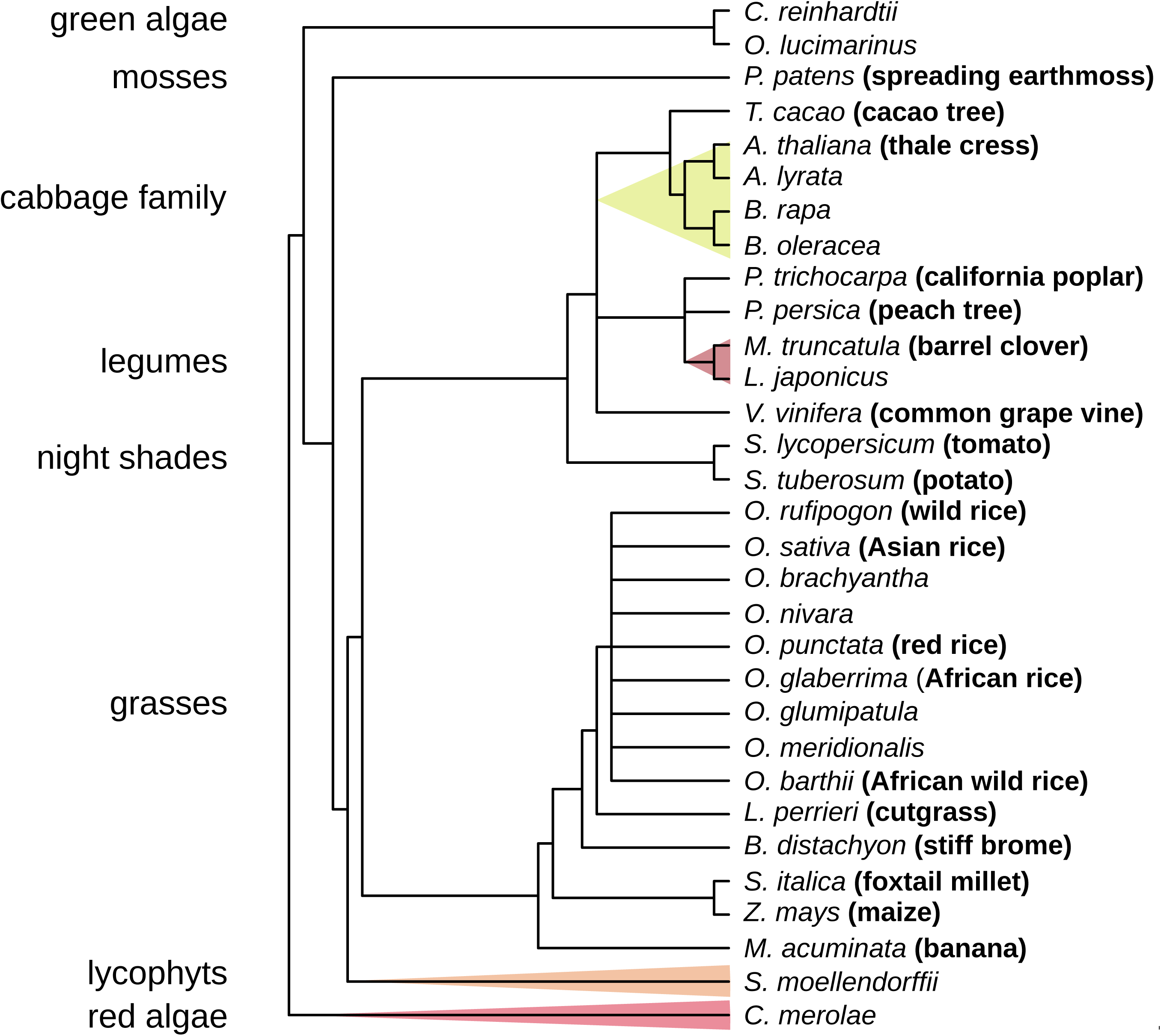
Taxonomic tree of the 31 investigated plant species. The tree is based on NCBI taxonomy and highlights some of the investigated plant families and their taxonomic relationship.

In our analysis, we used an artifice to detect putative new cluster types. Initially, the genomes under investigation were scanned for co-localized marker enzyme combinations of any two enzyme families. Additionally, we took advantage of the greediness of the search algorithm that automatically merges any marker enzymes detected within the cluster range and the flanking region with the cluster itself, as well as neighboring clusters if they are in close proximity (see Figure 1 for a graphical illustration). Using this artifice we were able to detect not only known MGCs but also novel combinations of marker enzymes that represent putative new cluster types. As additional criteria, in our further analysis we only considered those clusters that contained at least three marker enzymes of which at least two belonged to different enzyme families. Note, that the large superfamily of CYPs represents a collection of CYP sub-families which should be regarded as different types of marker enzymes but are represented by the same homolog PFAM domain and therefore cannot be distinguished by the search algorithm. To investigate the robustness of the search we performed the analysis for cluster ranges of 10 to 100 kb in-between each two enzymes and allowed for a 20kb flanking region (see Materials and Methods). Across this range, we obtained qualitatively similar results, more precisely; we obtained identical results for cluster ranges between 10 and 40 kb. Therefore, results from the 20kb cluster range sereach will be further analyzed and discussed in the following.

Our exhaustive search for co-localized enzymes from 26 specialized metabolism enzyme families resulted in the detection of in total 1232 putative MGCs types and a total number of 5531 putative clusters across the 31 plant species under investigation. The detected MGCs contained between 3 to 30 marker enzymes, whereas the most common cluster length across all species investigated was 3 (Figure 4A). Note, that throughout the analysis, gene clusters were considered as belonging to the same type of cluster, if their marker enzymes and the order along the chromosome were identical. This requirement was applied for the following two reasons; (i) in gene clusters that were characterized in several species the order of marker enzymes was typically (partially) conserved (8, 27, 28) and (ii) it was shown that genes in the noscapine MGC appeared to exhibit temporal collinearity, *i.e.* the gene order on the chromosome corresponds roughly to the temporal order of gene activation in the pathway (7). While this is the first example of this phenomenon in plants, collinearity was repeatability observed in bacteria and filamentous fungi and it might be observed in plants more often in future studies.

**Figure 4:**
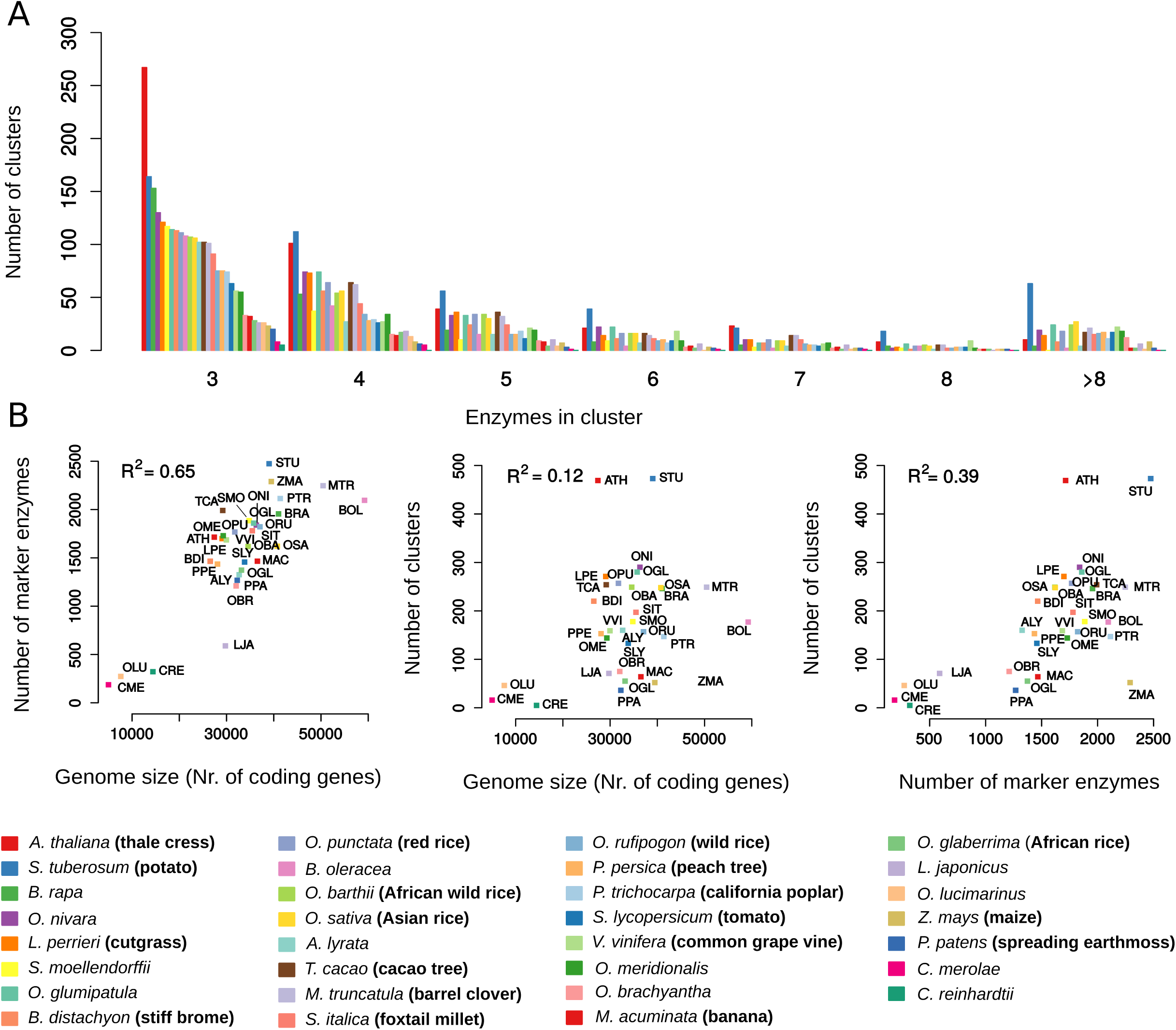
Characteristics of the detected MGCs --- Cluster size (number of marker enzymes) and relationship to genome size. A) Distribution of MGC sizes (number of marker enzymes) for all 31 investigated species. Shown is the absolute number of MGCs with different numbers of marker enzymes per species (common names are indicated in brackets if available). The most common cluster length is 3. B) Relationships between genome size, number of detected gene clusters, and number of members of secondary metabolism enzyme families (marker enzymes) for the 31 investigated species. Species names are indicated by a three-letter abbreviation code. Only the relationship between the number of marker enzymes and the genome size shows a positive correlation (R^2^=0.65) for the 31 investigated species.

To deepen our knowledge regarding the distribution and organization of individual marker enzymes across the genome, we performed a search for individual members of the 26 enzyme families across all genomes and compared it to the results of our MGC search. We found between 4.6 % (*Z. Mays*) to 57.1 % (Arabidopsis) of all detected marker enzymes to be organized in putative MGCs (mean over all species was 23.7%). As a comparison, Chae *et al.,* (2014) found that in Arabidopsis 30.1%, in soybean 30.2%, in sorghum 30.5%, and in rice 22.4% of genes are located in MGCs. Moreover, we observed a weak correlation between the overall number of marker enzymes and the genome size (R^2^ = 0.65) (Figure 4B, panel 1). Nevertheless, we did not observe significant correlations between the number of MGCs and genome size (R^2^ = 0.12) nor between the number of MGCs and the overall number of marker enzymes (R^2^ = 0.39) (Figure 4B, panel 2 and 3, respectively).

### Plant taxonomy is reflected by the types of MGCs present in the genomes

We subsequently tested whether the taxonomy of the investigated plant species was reflected by differences in the detected MGC types. First, we investigated similarities between species with respect to the detected cluster types. Clustering by MGC type similarity (see Materials and Methods) clearly reflected the taxonomy of the species under investigation as could be observed for algae, grasses, the nightshades, the single moss species, the single lycophyte, and members of the *Brassicaceae* (cabbage) family (Figure 5).

**Figure 5:**
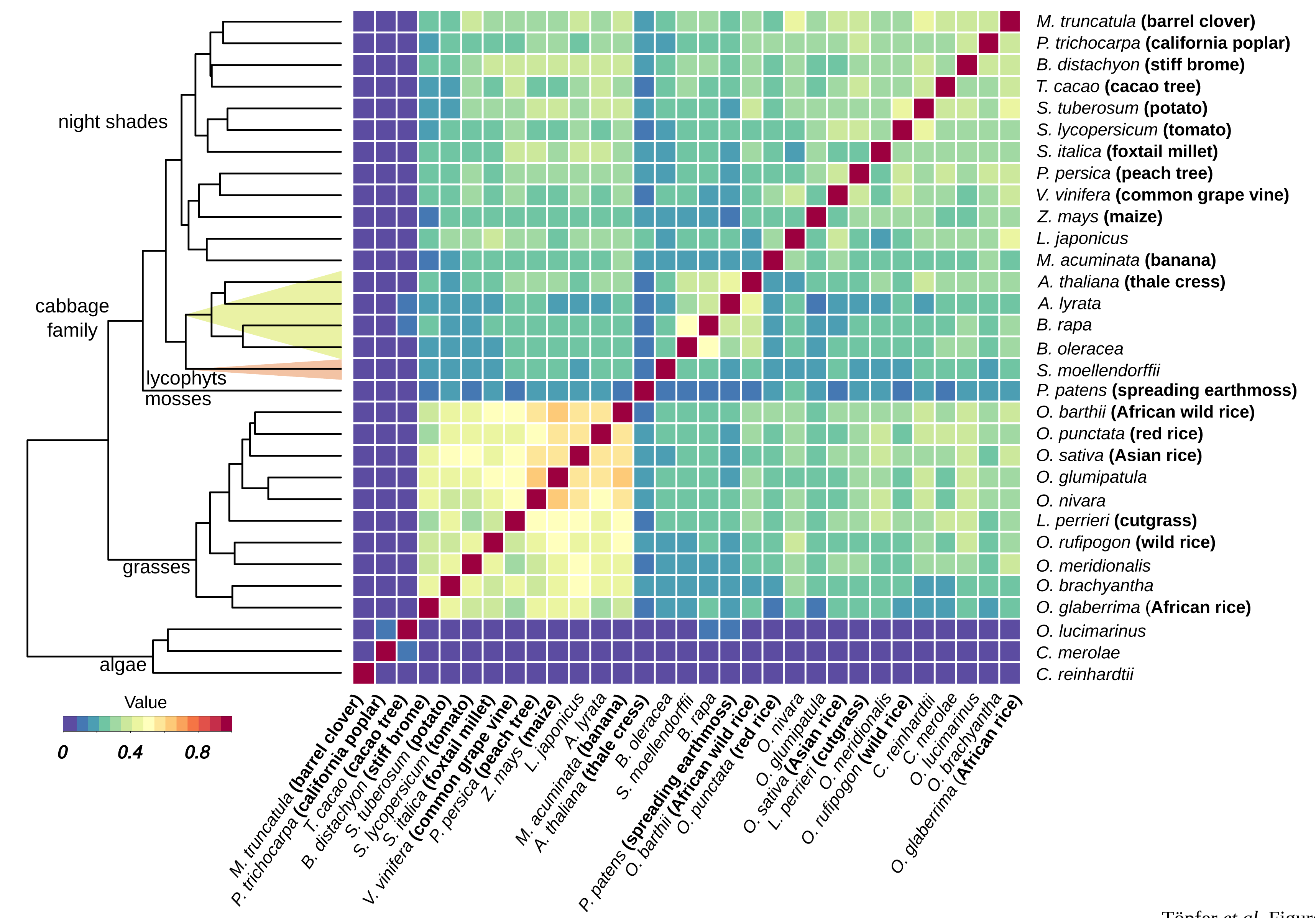
Green lineage taxonomy is reflected by MGCs. Clustering analysis of the 31 investigated species genome based on MGCs similarity. The cosine similarity based on the presence or absence of MGC types was used as a similarity measure to calculate the clustering profile. Related species cluster together, such as algae, grasses, the nightshades, the single moss species, the single lycophyte, and members of the *Brassicaceae* (cabbage) family (compare to taxonomy in Figure 3).

### Enzyme families significantly enriched in MGCs

We next investigated which enzyme families are enriched in MGCs and whether taxa-specific differences could be found. When performing enrichment analysis (Fisher’s-exact test) across all species we found 2-oxoglutarate-dependent dioxygenases (*p* = 7.2e-31), CYPs (*p* = 1.3e-02), glutathione-*S*-transferases (*p* = 1.9e-88), methylene bridge-forming enzymes (*p* = 6.6e-22), terpene synthases (*p* = 4.7e-62), and 2-Isopropylmalate synthases (*p* = 7.3e-10) to be significantly enriched in MGCs across the plant kingdom. Notably, when performing the same analysis with a relaxed cluster definition (including clusters with only two marker genes and tandem repeats) we found all enzyme families but CYPs (*p*=1.03e-164) to be less significant (Supplementary Information 3). This observation supports the notion of an accelerated rate of gene duplications in certain CYPs clans (29) which are likely to make up most of the tandem repeat gene clusters which are not excluded from the analysis when using the more relaxed constraints.

Taxa-specific analysis revealed notable differences between taxonomic groups *e.g.,* members of the glutathione-*S*-transferases family were found enriched in MGCs across the plant kingdom, whereas 2-oxoglutarate-dependent dioxygenases and methylene bridge-forming enzymes were enriched in angiosperms only and CYPs in monocots only. An overview of significantly enriched enzyme families in MGC across all investigated species and for algae, mosses, lycophytes, mono- and dicots separately is shown in Figure 6. Figure 7 shows which enzyme families are enriched in MGCs in each of the investigated species (see also Supplementary Information 4 for analysis of enriched enzyme combinations).

**Figure 6:**
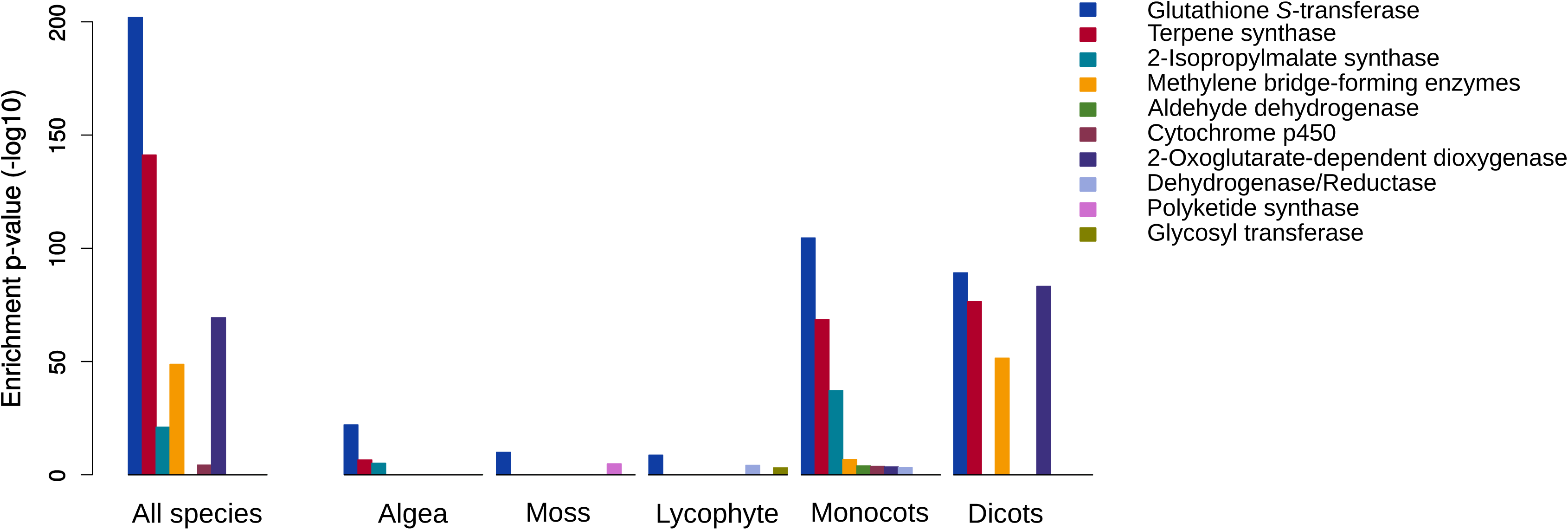
Enrichment of marker enzymes in putative MGCs in different phylogenetic taxa. A) Enrichment analysis for specialized metabolism associated enzyme families was performed (i) for all species together (left side) and (ii) the five investigated taxa (algae, mosses, lycophytes, monocots, and dicots) separately (right side). The y-axis represents the negative log_10_ transformation of the *p-*value for the enrichment based on the Fisher’s exact test. Shown are only significant enzyme families (*p*-value < 0.05). Members of the glutathione-*S*-transferases family were found enriched in MGCs across the plant kingdom, whereas 2-oxoglutarate-dependent dioxygenases and methylene bridge-forming enzymes were enriched in angiosperms only and CYPs in monocots only.

**Figure 7:**
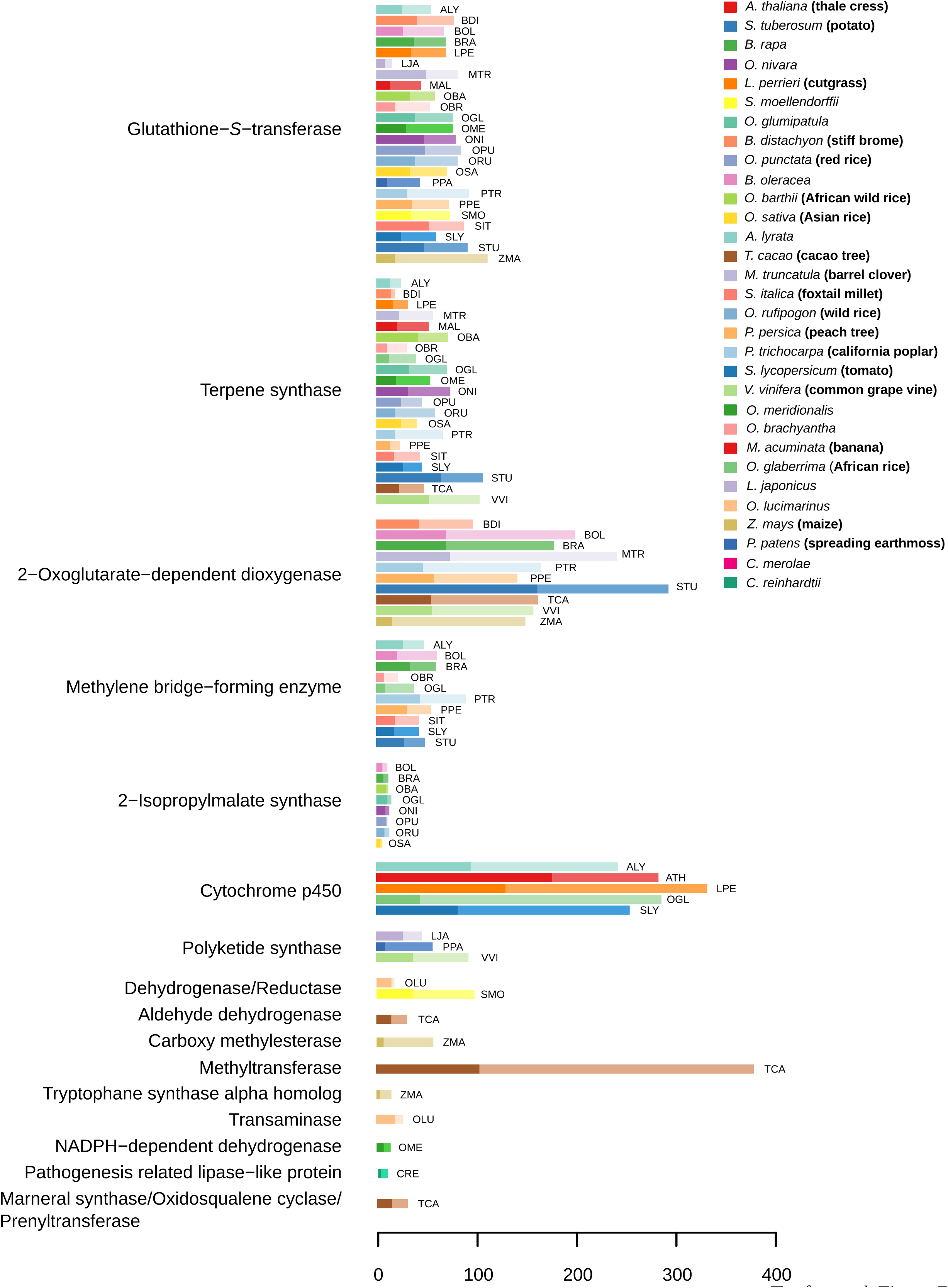
Enrichment of marker enzymes in putative MGCs in the 31 investigated species genomes. The bar plots show the total number of marker enzymes (lighter color) and the number of marker enzymes organized in putative MGCs (darker color) for all marker enzyme families and all species separately. Shown are only enzyme families and species which exhibit a significant enrichment of the respective marker enzymes in putative MGCs based on Fisher’s exact test (*p*-value < 0.05). Species names are indicated by a three-letter abbreviation code (common names indicated in brackets).

### Combining MGCs prediction with gene co-expression analysis results in the identification of likely candidates

We performed co-expression analysis for Arabidopsis, rice, and tomato to investigate which MGCs contain significantly co-expressed genes with respect to the reference background co-expression. As a reference we used the co-expression patterns of all marker genes in the respective genome and examined (i) the mean correlation of marker genes in MGCs against the background distribution, and (ii) the correlation distribution of marker genes in individual MGCs against the background distribution (see Materials and Methods for details). We found statistically significant higher co-expression between marker genes in MGCs with respect to the background distribution only for tomato (Wilcoxon Rank test; *p* = 0.001). Nevertheless, when investigating the correlation distribution of individual MGCs against the background we found genes in 4 clusters in Arabidopsis, 3 in rice, and 7 in tomato to be significantly higher co-expressed (Wilcoxon rank test, *p* < 0.05). The length of these clusters varied between 51 and 140 kb and they contained between 4 and 18 marker enzymes. The results of this analysis are summarized in Figure 8. A complete list of the detected marker enzymes and the respective genes are given in the Supplementary Information 5. Interestingly, when running the same analysis for the relaxed cluster definition (including also clusters with only two marker genes and tandem repeats) we found increased and statistically significant co-expression of marker genes in the MGCs for all three species examined (Wilcoxon Rank test, *p* = 0.002, 3.04e-4 and 2.8e-11 for Arabidopsis, rice and tomato, respectively).

**Figure 8:**
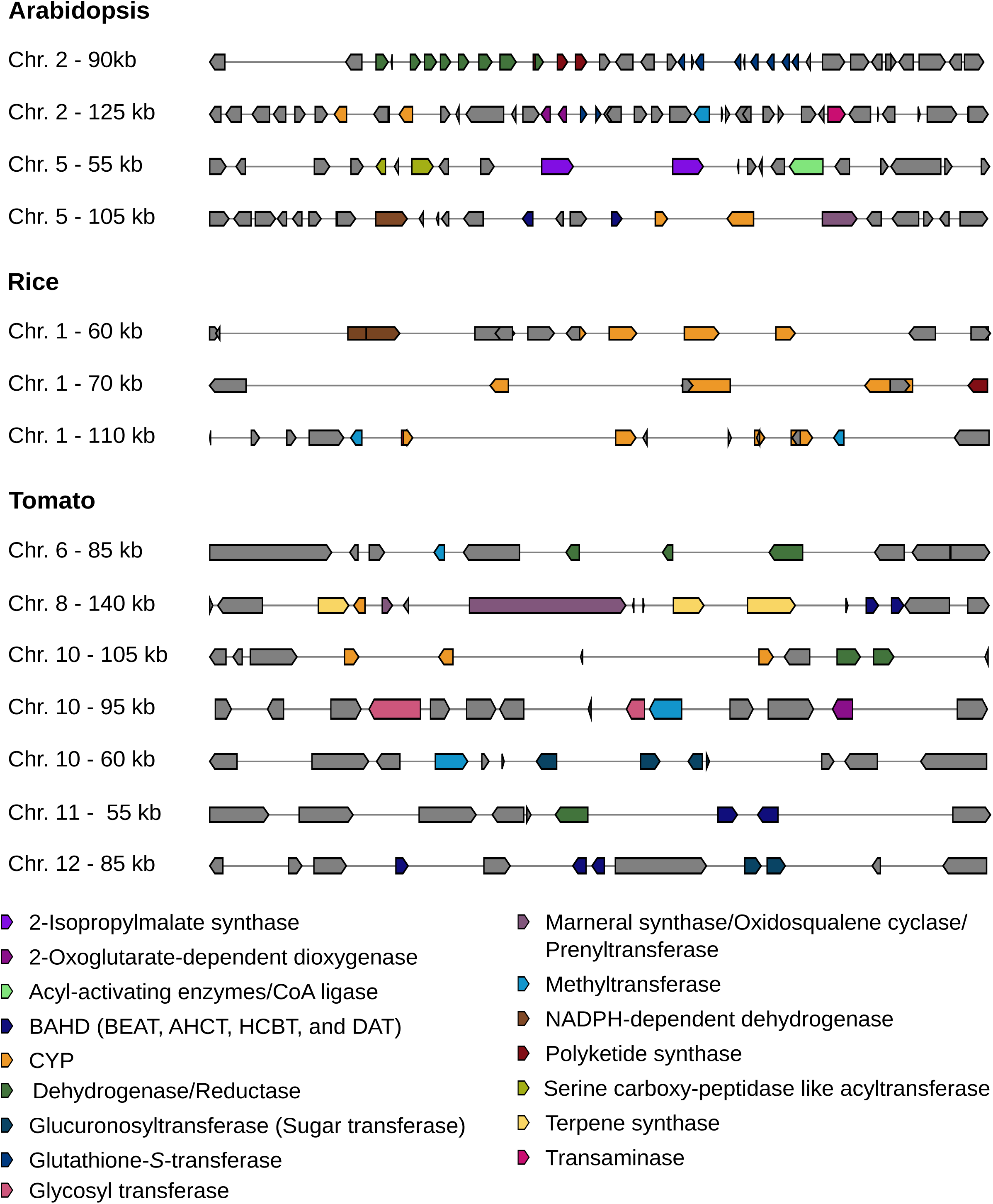
Putative MGCs with statistically significant gene co-expression. The schematic overview shows the candidate clusters that are significantly higher co-expressed with respect to the background distribution including all detected marker enzymes (Wilcoxon Rank test, *p* < 0.05). The left side shows the chromosomal location and length of the putative cluster and the right side depicts the organization of the marker enzymes along the chromosome.

## DISCUSSION

In this study we developed a computational tool for the identification and analysis of putative MGCs in plants. The tool accurately detects known MGCs. An exhaustive search of 31 genome assemblies across the plant kingdom resulted in the detection of several thousand MGCs candidates. Furthermore, we found genes associated with specialized metabolism enzymes to exhibit taxa-specific co-location patterns, reflecting the underlying taxonomy of the species. Enrichment analysis revealed taxa- and species-specific enrichment of certain enzyme families. Co-expression analysis for Arabidopsis, rice, and tomato revealed an increased level of co-expression of cluster candidates only in tomato. Nevertheless, statistical analysis of individual gene cluster candidates yielded a list of putative MGCs containing co-expressed genes of statistical significance in Arabidopsis, rice, and tomato.

### The discovery of additional plant MGCs is crucial for refining cluster detection rules

PhytoClust represents a highly exhaustive search algorithm as any combination of specialized metabolism enzyme classes can be detected. Conversely, the “known-cluster search algorithm” in PhytoClust is specific and detects only complete MGCs of a particular enzyme class composition. Therefore, evolutionary modifications of the same cluster in different species (*e.g.,* loss of a certain marker enzyme or splitting of clusters) would not necessarily be detected. Future insights into the organization and evolution of MGCs in plants as well as deeper knowledge of metabolic pathways will help reducing the gap between these two approaches. As a result, cluster rules in PhytoClust could be attuned according to *e.g.,* certain chemical classes produced by particular combinations of enzymes while being flexible with respect to additional marker enzymes in the cluster.

Our study points at the importance of generating specific HMMs for members of the CYP family. More than 5100 sequences of the plant CYP family were described which are likely involved in numerous functions. For the larger families within the clan, enzymes with even less than 40 % sequence similarity are grouped into sub-families (29). Despite this tremendous diversity, currently all CYPs have the same HMM in the PFAM database (30). A more sophisticated representation of these enzyme families would significantly improve the predictive power of the PhytoClust search algorithm. Likewise, we anticipate the repertoire of marker enzymes to expand as new MGCs will be discovered and other classes of gene products, such as enzymes involved in primary metabolism, transcription factors, signaling components or transporters, will be found as MGCs components.

Moreover, any analysis of putative MGCs, *e.g.,* enrichment of certain enzyme families in MGCs, heavily depends on the chosen set of putative MGCs. Relaxation of the cluster search definition from three to two marker enzymes and allowance for tandem repeats resulted in (i) higher over-representation of members of the CYPs class and (ii) rendered enzymes from putative MGCs to be significantly co-expressed in the case-study of Arabidopsis, rice, and tomato. This might explain different findings with respect to previous studies (10). Moreover, our findings emphasize the significant role of CYPs in specialized metabolism gene duplication and neo-functionalization that drives gene diversification and potentially MGCs evolution.

### The potential of plant MGCs for rational design strategies

The discoveries of MGCs in plants through programs such as PhytoClust will likely speed-up metabolic pathway elucidation. Yet, it is also expected to provide the information required for subsequent metabolic engineering of pathways generating high-value products in plants or microorganisms (5). Already at this stage where a relatively small number of MGCs has been characterized in depth, MGCs associated with the biosynthesis of high-value molecules have been discovered *e.g.,* noscapine, an antitumor alkaloid from opium poppy (7). Moreover, MGCs provide a natural example for pathway engineering and could therefore teach us how to tackle issues of coinheritance, avoidance of toxic intermediates, spatial and temporal control of gene expression, metabolic channeling and likely much more. Hence, we expect that working with PhytoClust will boost the elucidation of metabolic pathways of specialized metabolism, understanding their evolution and engineering. These advances are nevertheless largely dependent on the quality and number of new plant genomes assembled in the coming years.

## MATERIALS AND METHODS

### Translating specialized metabolism enzyme families into hidden Markov models

We compiled a list of 26 enzyme families known to catalyze reactions in plant specialized metabolism (*i.e.* “marker enzymes”) through literature mining (4, 31–33). This manually curated list was subsequently translated into a library of hidden Markov models by extracting relevant entries from the PFAM database (30) (Supplementary Information 1). Using this mathematical representation of protein families we were able to represent combinations of marker enzymes by cluster rules. These cluster rules in turn are used to search the genome of interest for combinations of these enzymes within a given distance (“cluster range”). Following the AntiSmash core implementation (18), the flanking regions of a detected cluster are extended by 20 kb on each side of the outer marker enzymes. All marker enzymes from the library that are detected within this extended cluster region are listed in the results. As a consequence of this procedure, the search algorithm is rendered comprehensive and enables the detection of novel MGCs types that were not explicitly searched for.

### Input genomes and databases

A collection of publicly available plant genomes was subjected to analysis by PhytoClust. For reasons of reproducibility and runtime constraints we chose a selection of genomes with the highest assembly quality that required a relatively short runtime. Calculations on a 64bit Linux computer with 3.00GHz × 16 took between minutes to days depending on genome size and quality as well as on the applied cluster rules. All genomes (except for *Lotus japonicus*) used for this analysis were downloaded as .embl files from Ensemble plants release 31 (34). A comparative analysis with genomes downloaded previously from Ensemble plants release 25 resulted in the same outcome. The genome for *L. japonicus* was obtained from the PlantGDB (Version 1.0) (35).

### Sensitivity of MGC prediction to different proximity ranges

We performed a sensitivity analysis for our MGC search algorithm by re-running the algorithm for 10, 20, 30, 40, 60, 80, 100kb proximity of any combination of two marker enzymes. Significantly, the results did not change for proximity ranges between 10 and 40 kb. Therefore, we chose the combination of 20kb cluster range and 20kb flanking region as the default settings for the presented analysis. Across all investigated proximity ranges the total number of detected clusters varied between 5355 and 5694. Note that for different cluster ranges, the detected MGCs do not necessarily need to be identical. Differences may arise as marker enzymes in the vicinity of the cluster might be detected and included in the putative MGC or not depending on the chosen proximity range.

Note that the used cluster range in our search for combinations of any two enzymes is not equal to the length of the detected cluster as a whole but rather represents the maximum distance between two adjacent marker enzymes. Therefore, depending on the number of marker enzymes in the cluster the complete cluster can be longer.

### Co-expression module

The co-expression module is currently available for *A. thaliana*, *S. lycopersicum*, *S. tuberosum,* and *O. sativa* based on publically available datasets (8, 36–38). The algorithm performs Pearson-correlation based co-expression analysis and plots a heat-map for those genes which are (i) present in the transcript data and (ii) co-expressed under these particular experimental conditions, *i.e.* these results can only be treated as a support for the identification of a MGC candidate but not as a dissaproval. A gene cluster that does not show co-expression in our pipeline might as well exhibit coordinated expression when using either a more comprehensive dataset or data from different experimental conditions.

### Statistical analysis

Statistical analyses were performed using R and the “stats” package for Fisher Exact test (fisher.test), Wilcoxon Rank test (wilcox.test), and the “Isa” package for Cosine similarity (cosine). The cluster similarity heat-map was generated using the “gplots” package. As input for the cosine matrix calculation we used a binary matrix of ones and zeros, indicating whether a certain cluster type was detected in the species or not.

## ACKNOWLEDGEMENT

We thank the Adelis Foundation, Leona M. and Harry B. Helmsley Charitable Trust, Jeanne and Joseph Nissim Foundation for Life Sciences, Tom and Sondra Rykoff Family Foundation Research and the Raymond Burton Plant Genome Research Fund for supporting the AA lab activity. AA is the incumbent of the Peter J. Cohn Professorial Chair. We thank Ronen Levi and Gabor Szabo for logistic support.

### FUNDING

N.T. was supported through the Deans of Life Sciences Postdoctoral fellowship and the Alternative Energy Research Initiative (AERI) of the Weizmann Institute.

